# A Ca^2+^ wave generates a force during cell extrusion in zebrafish

**DOI:** 10.1101/2021.07.14.452309

**Authors:** Sohei Yamada, Yasumasa Bessho, Yasuyuki Fujita, Yoichiroh Hosokawa, Takaaki Matsui

**Affiliations:** Division of Materials Science, Nara Institute of Science and Technology, 8916-5 Takayama, Ikoma, Nara 630-0192, Japan; Graduate School of Science and Technology, Hirosaki University, 3 Bunkyo-Cho, Hirosaki, Aomori, 036-8561, Japan; Division of Biological Science, Nara Institute of Science and Technology, 8916-5 Takayama, Ikoma, Nara 630-0192, Japan; Department of Molecular Oncology, Graduate School of Medicine, Kyoto University, Kyoto, 606-8501, Japan

## Abstract

When oncogenic transformed or damaged cells appear within an epithelial sheet, they are apically extruded by surrounding cells. Recently, using cultured mammalian epithelial cells and zebrafish embryonic epithelial cells, we found that a calcium (Ca^2+^) wave propagates from RasV12-transformed cells and laser-irradiated damaged cells to surrounding cells and promotes apical extrusion by inducing polarized movements of the surrounding cells. In mammalian cell cultures, we reported that the inositol trisphosphate (IP_3_) receptor, gap junctions, and the mechanosensitive Ca^2+^ channel TRPC1 are involved in Ca^2+^ wave-mediated polarized movements. However, which molecules regulate Ca^2+^ wave-mediated polarized movements in zebrafish and whether the Ca^2+^ wave can generate a force remain unknown. In this study, we aimed to answer these questions. By performing pharmacological and gene knockout experiments, we showed that a Ca^2+^ wave induced by the IP_3_ receptor and trpc1 led to formation of cryptic-lamellipodia and polarized movements of surrounding cells toward extruding cells in zebrafish. By using an *in vivo* force measurement method, we found that the Ca^2+^ wave generated approximately 1 kPa of force toward extruding cells. Our results reveal a previously unidentified molecular mechanism underlying the Ca^2+^ wave in zebrafish and demonstrate that the Ca^2+^ wave generates a force during cell extrusion.

## Introduction

Epithelial sheets act as a barrier that maintains tissue integrity and homeostasis. When oncogenic transformed or damaged cells appear in these sheets, they induce loss of tissue integrity and can lead to microbial infection and tumorigenesis (*1, 2*). Thus, multicellular organisms have developed a system that eliminates these cells from the sheets without disrupting epithelial integrity via a process called cell extrusion (*3–5*).

It has been proposed that actomyosin ring contraction generates a driving force for cell extrusion (*6*). When oncogenic transformed or damaged cells (extruding cells) appear in epithelial sheets, filamentous actin (F-actin) polymerizes at the adjacent edge of each cell surrounding the extruding cells. Subsequently, myosin-2 is activated by the RhoA/ROCK pathway, leading to formation of actomyosin bundles in each surrounding cell (*5,7,8*). Thereafter, the actomyosin bundles are coupled with intercellular adhesions to form an actomyosin ring between surrounding cells. Finally, the actomyosin ring shrinks at the basal level and pushes extruding cells outward apically. To determine whether actomyosin ring contraction generates the force required for cell extrusion, we previously developed a method using a femtosecond laser-induced impulsive force and revealed that the contractile force of the actomyosin ring is approximately 4 kPa in zebrafish, indicating that actomyosin ring contraction is one of the force-generating processes during cell extrusion (*9*).

Recently, using mammalian cell cultures and zebrafish, we found that a Ca^2+^ wave propagates from extruding cells toward surrounding cells before actomyosin ring contraction (*10*). The Ca^2+^ wave induces polarized movements of surrounding cells toward extruding cells, and initiates apical extrusion, at least in part, by inducing actin rearrangements in surrounding cells. Furthermore, in mammalian cell cultures, the inositol trisphosphate (IP_3_) receptor, gap junctions, and the mechanosensitive Ca^2+^ channel TRPC1 are involved in Ca^2+^ wave-mediated polarized movements. However, the molecular mechanism underlying Ca^2+^ wave-mediated polarized movements in zebrafish and whether the Ca^2+^ wave can generate a force required for cell extrusion remain unknown.

In this study, we demonstrated that the Ca^2+^ wave regulates the formation of lamellipodia in multiple rows of surrounding cells, so-called cryptic-lamellipodia (c-lamellipodia), which regulate the polarized movements of collective cell groups (*11–14*). At the molecular level, we showed that the IP_3_ receptor and mechanosensitive Ca^2+^ channel TRPC1 are involved in Ca^2+^ wave-mediated polarized movements. By using the *in vivo* force measurement method that we previously established, we found that the extruding force generated by the Ca^2+^ wave is approximately 1 kPa. Based on the results of our previous and current studies, we propose that cell extrusion is driven by relatively small forces (1–4 kPa) generated by surrounding cells and that at least two distinct processes (i.e., Ca^2+^ wave-mediated polarized movements and actomyosin ring contraction) contribute to force generation during cell extrusion.

## Results

### Ca^2+^ wave propagation during cell extrusion in zebrafish

We previously established a method to induce cell extrusion in zebrafish embryos. When the center of an epithelial cell is irradiated with a femtosecond laser in zebrafish embryos, the irradiated damaged cell is apically extruded by surrounding cells within 5 min (*9, 10*). To investigate Ca^2+^ and F-actin dynamics during cell extrusion in zebrafish, we applied this method to embryos expressing GCaMP7 (a Ca^2+^ sensor) (*15*) and Lifeact-GFP (a marker of F-actin) (*16*). Immediately after laser irradiation, Ca^2+^ levels increased in irradiated (extruding) cells, and then a Ca^2+^ wave propagated from the extruding cells to surrounding non-irradiated cells (Fig. 1A, B). Ca^2+^ levels in the surrounding cells peaked within 60 s after irradiation and decreased to the basal levels by 100 s (Fig. 1A, B, Video S1). As shown recently, the Ca^2+^ wave could propagate approximately 2–7 cell lengths (*10*) (see also Fig. 1A, B). After Ca^2+^ wave propagation (more than 120 s after irradiation), the actomyosin ring, as evidenced by the distribution of F-actin, formed between surrounding cells adjacent to the extruding cells, indicating that the Ca^2+^ wave propagated before actomyosin ring contraction (Fig. 1A) (*10*).

**Figure 1.**
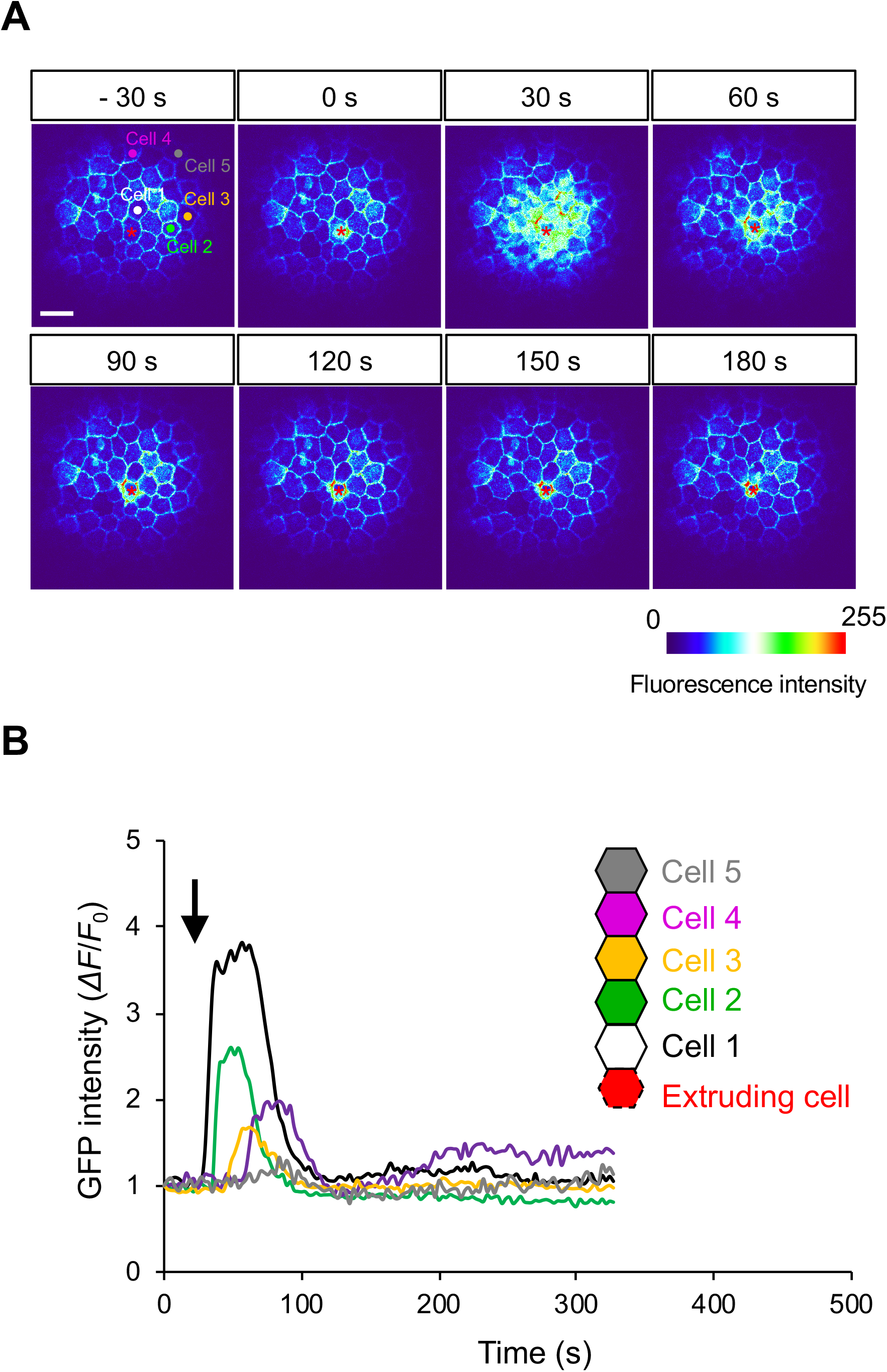
Ca^2+^ wave propagation during cell extrusion in zebrafish. (A) Color-coded images of GCaMP7 and Lifeact-GFP dynamics during cell extrusion. Still images were extracted from Video S1. Red asterisks indicate the laser focal point. Scale bar: 50 μm. (B) The fluorescence intensity of GCaMP7 was quantified in multiple rows of surrounding cells around the extruding cell. White, green, yellow, purple, and gray lines show Ca^2+^ levels in the first, second, third, fourth, and fifth rows of surrounding cells, respectively, as shown in (A). We calculated *ΔF/F*_0_, where *ΔF* is the instantaneous fluorescence signal and *F*_0_ is the fluorescence intensity in the initial frame.

### Ca^2+^ wave-mediated polarized movements and actin rearrangements in zebrafish

During apical extrusion, surrounding cells must fill the vacant space. We recently demonstrated that during cell extrusion in mammalian cell cultures, the Ca^2+^ wave induces F-actin accumulation in the cytosol and perinuclear regions of surrounding cells, leading to orchestrated movements of surrounding cells toward extruding cells (*10*). However, we did not characterize the cellular mechanisms underlying Ca^2+^ wave-mediated polarized movements during cell extrusion in zebrafish. Thus, we investigated these mechanisms in zebrafish. By analyzing the movement of vertices of surrounding cells located inside or outside the Ca^2+^ wave (Fig. 2A) (*10*), we found that vertices inside the Ca^2+^ wave moved further than those outside the Ca^2+^ wave (Fig. 2B). Furthermore, surrounding cells inside the Ca^2+^ wave preferentially moved toward the extruding cells, but those outside the Ca^2+^ wave did not exhibit such polarized movements (Fig. 2B, C). These phenotypes are consistent with those of mammalian cultured cells (*10*). Although the polarized movements of surrounding cells continue until the completion of cell extrusion in mammalian cell cultures, they were only observed during Ca^2+^ wave propagation in zebrafish (Fig. 2B).

**Figure 2.**
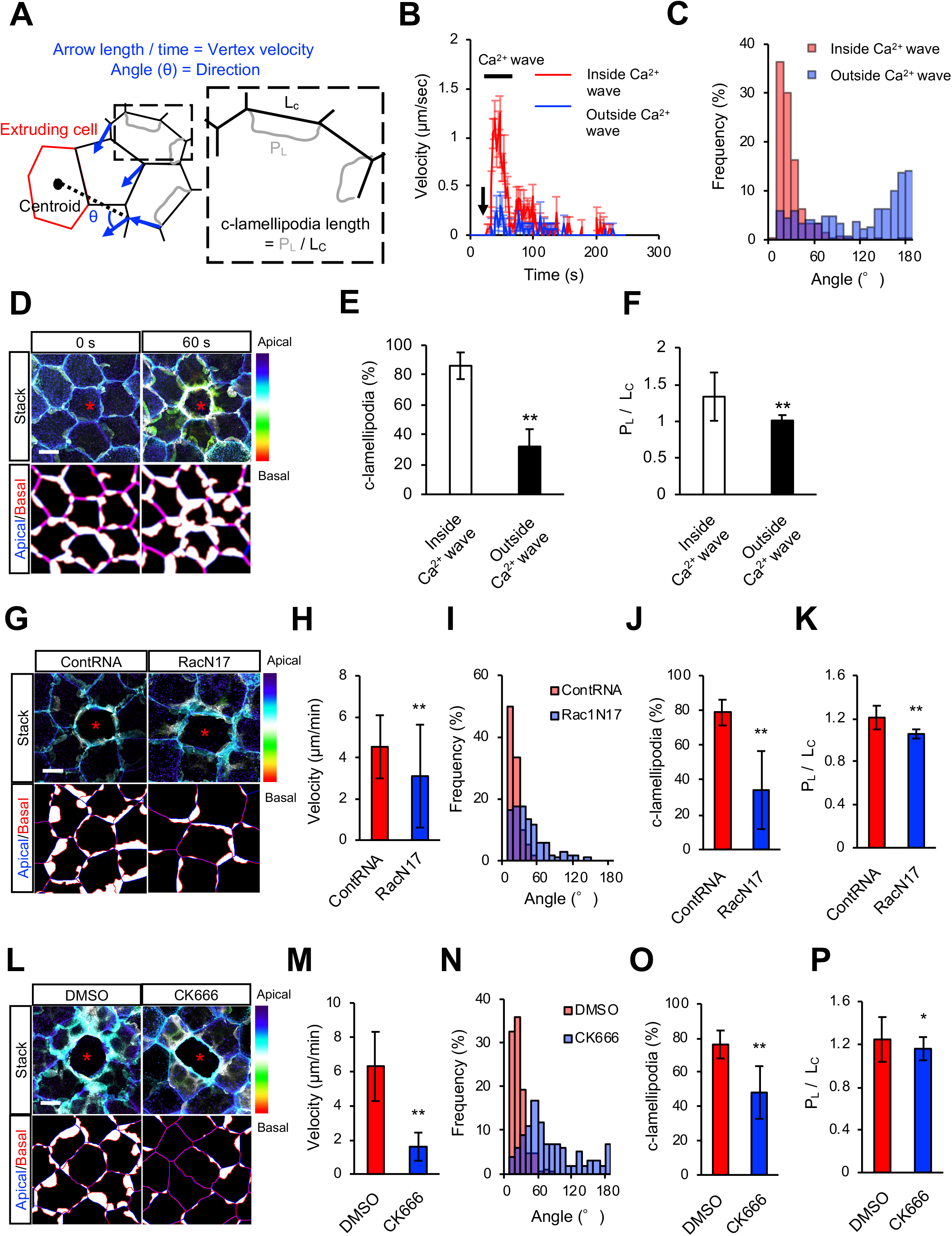
c-lamellipodia form in surrounding cells inside the Ca^2+^ wave. (A) Left panel: schematic diagram of the displacement and direction of movement of cell vertices during cell extrusion. Blue arrows denote vertex movement from start to end. The arrow angle (θ) forms between the end and start vertex and the centroid of the extruding cell. The arrow length and angle indicate the displacement and direction of the vertex movement, respectively. Right panel: schematic diagram of the formation and elongation of c-lamellipodia. To estimate c-lamellipodia formation, we measured the length of lamellipodia (P_L_)and contact length (L_C_) and calculated P_L_/L_C_. (B) The velocity of vertex displacement of surrounding cells inside (red, n = 12) and outside (blue, n = 11) the Ca^2+^ wave. The black arrow and bar indicate the time of laser irradiation and duration of the Ca^2+^ wave, respectively. (C) Quantification of the direction of vertex movement. Red and blue histograms show the direction of surrounding cells inside (red) and outside (blue) the Ca^2+^ wave. n = 314 and 316 from 15 embryos. (D) Distribution of F-actin in the different apical-basal z-planes at 0 and 60 s after laser irradiation. Upper panels, projections of apical-basal z-planes with blue-to-red colors as indicated to the right. Red asterisks indicate the laser focal point. Scale bar: 10 μm. Lower panels, schematic illustrations of F-actin distributions, projected with the apical (top) and basal (bottom) z-planes. The white area indicates c-lamellipodia. (E-F) Frequency (E) and size (F) of c-lamellipodia in surrounding cells inside (white) and outside (black) the Ca^2+^ wave. n = 316 and 256 cells from nine embryos. (G–P) Effects of RacN17 overexpression (G–K) and CK666 treatment (L–P) on polarized movements and c-lamellipodia formation. Representative images of the F-actin distribution at 60 s after laser irradiation (G, L). In control embryos (contRNA-injected or DMSO-treated), polarized movements (H, I, M, N, red) and formation of c-lamellipodia (J, K, O, P, red) occurred normally. However, in embryos injected with RacN17 mRNA (G–K) or treated with CK666 (L–P), polarized movements (H, I, M, N, blue) and formation of c-lamellipodia (J, K, O, P, blue) were inhibited. (H) n = 39 and 56 cells from three and five embryos, respectively. (I) n = 60 and 102 from three and five embryos, respectively. (J and K) n = 21 and 36 cells from four and six embryos, respectively. (M) n = 58 and 81 cells from six and five independent experiments, respectively. (N) n = 183 and 112 from six and five embryos, respectively. (O and P) n = 21 and 36 cells from four and six embryos, respectively. Error bars show SD. \**P* < 0.05; \*\**P* < 0.01 (paired t-test).

We next investigated F-actin dynamics during the Ca^2+^ wave in zebrafish. F-actin accumulated at lamellipodia in multiple rows of surrounding cells, which exhibited front-to-rear polarity toward extruding cells (Fig. 2D). Their structures are similar to those of c-lamellipodia required for collective migration of cohesive cell groups during wound healing (*13, 14*). It has been reported that Rac1 regulates Arp2/3-mediated actin polymerization, leading to formation of c-lamellipodia (*12, 14*). To investigate whether the F-actin-enriched lamellipodia that formed in multiple rows of surrounding cells are c-lamellipodia, we inhibited the Rac1-Arp2/3 pathway in our experimental settings. Overexpression of RacN17 (dominant-negative mutant) or treatment with CK666 (Arp2/3 inhibitor) significantly reduced the formation rate and size of lamellipodia in surrounding cells and inhibited the polarized movements and collective behaviors of surrounding cells (Fig. 2G–P), indicating that the Ca^2+^ wave induces formation of c-lamellipodia during cell extrusion in zebrafish. We next investigated the relationship between Ca^2+^ wave propagation and c-lamellipodia formation. The c-lamellipodia formed in surrounding cells both inside and outside the Ca^2+^ wave, but the size and frequency of c-lamellipodia were significantly higher inside the Ca^2+^ wave than outside the Ca^2+^ wave (Fig. 2E, F). Importantly, reduction of c-lamellipodia formation by inhibition of the Rac1-Arp2/3 pathway inhibited the polarized movements of surrounding cells (Fig. 2H, I, M, N). Our results suggest that the Ca^2+^ wave induces the polarized movements of surrounding cells through the formation of c-lamellipodia in zebrafish.

### The mechanosensitive Ca^2+^ channel trpc1 and IP_3_ receptor are involved in the Ca^2+^ wave in zebrafish

By performing pharmacological analyses and gene knockdown in mammalian cell cultures, we showed that the IP_3_ receptor, gap junctions, and the mechanosensitive Ca^2+^ channel TRPC1 induce Ca^2+^ wave propagation (*10*), whereas we did not characterize the molecular mechanism in zebrafish. Therefore, we tested whether similar molecular mechanisms regulate the Ca^2+^ wave in zebrafish by analyzing the effects of several inhibitors of Ca^2+^ channels. Treatment with gadolinium (Gd^3+^; mechanosensitive Ca^2+^ channel inhibitor), GsMTx4 (mechanosensitive Ca^2+^ channel inhibitor), xestospondin C (XestoC; IP_3_ receptor inhibitor), or 2-aminoethoxydiphenylborane (2-APB; mechanosensitive Ca^2+^ channel and IP_3_ receptor inhibitor) diminished Ca^2+^ wave propagation, but treatment with 18α-glycyrrhetinic acid (a GA; gap junction inhibitor) did not (Fig. 3A, B). These results suggest that mechanosensitive Ca^2+^ channels and IP_3_ receptors, but not gap junctions, regulate the Ca^2+^ wave in zebrafish. Gd^3+^, GsMTx4, and 2-APB inhibit various types of mechanosensitive Ca^2+^ channels, but they all target TRPC1 and TRPC6 channels (*10*). Importantly, by performing gene knockdown experiments, we reported that TRPC1, but not TRPC6, regulates Ca^2+^ wave propagation in mammalian cultured cells (*10*). Thus, we generated *trpc1*-knockout zebrafish using the CRISPR. Consistent with the results from mammalian cell cultures, the Ca^2+^ wave was impaired in heterozygous and homozygous *trpc1* mutants (Fig. 4A, B), indicating that TRPC1 is the crucial regulator of Ca^2+^ wave propagation in mammalian cell cultures and zebrafish.

**Figure 3.**
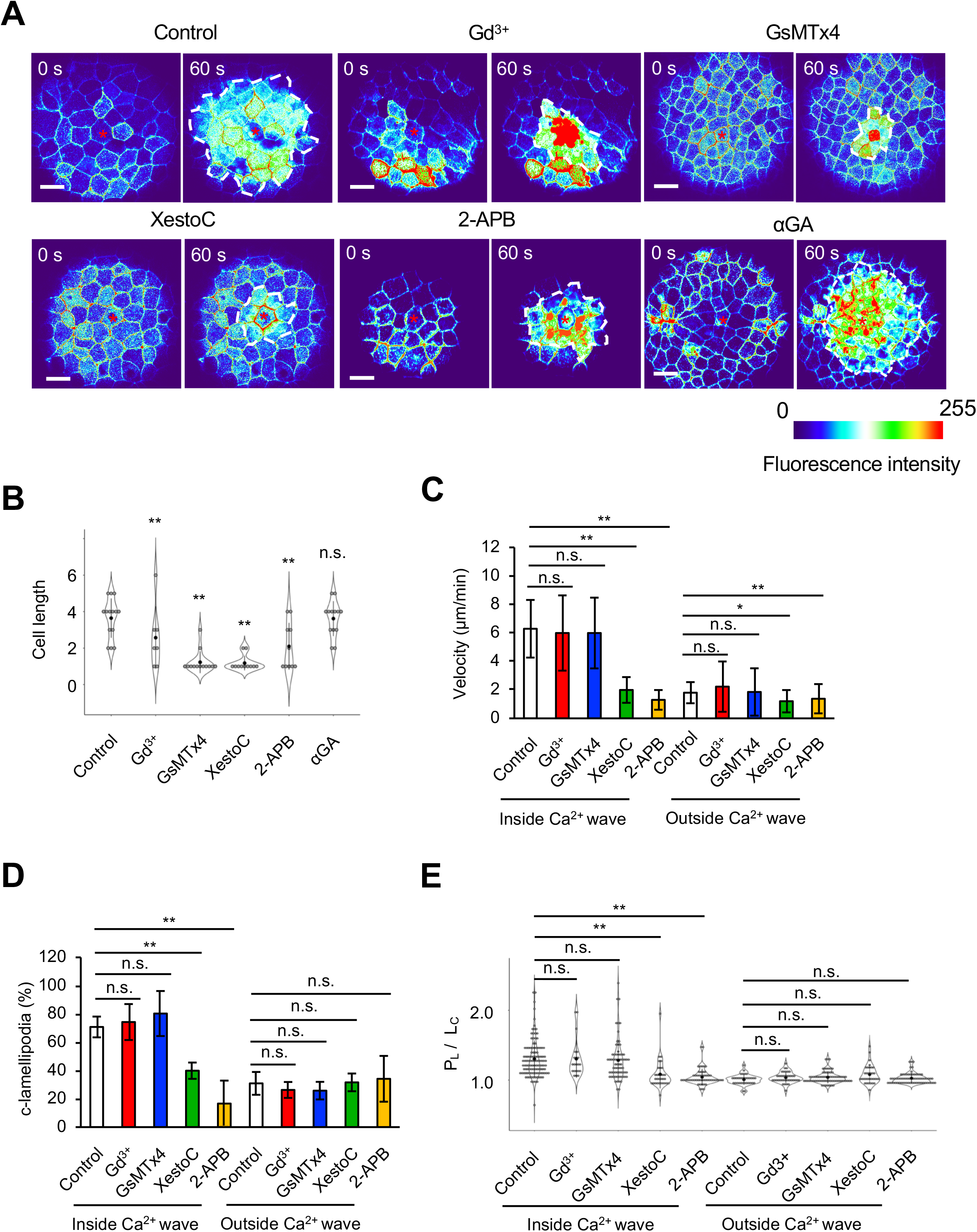
Mechanosensitive Ca^2+^ channels and the IP_3_ receptor are involved in Ca^2+^ wave-mediated polarized movements. (A) Color-coded images of GCaMP7 and Lifeact-GFP in embryos treated with DMSO (control), Gd^3+^, GSMTx4, XestoC, 2-APB, or αGA at 0 and 60 s after laser irradiation. White dotted lines indicate the area of Ca^2+^ wave propagation. Red asterisks indicate the laser focal point. Scale bar: 50 μm. (B–E) Effects of inhibitors on Ca^2+^ wave-mediated polarized movements. (B) The expansion area of the Ca^2+^ wave. n = 14, 7, 13, 12, 10, and 13. (C) Polarized movements of surrounding cells inside and outside the Ca^2+^ wave. n = 58 and 42 from eight embryos for control, n = 41 and 39 from five embryos for Gd^3+^ treatment, n = 58 and 58 from nine embryos for GsMTx4 treatment, n = 58 and 58 from seven embryos for XestoC treatment, and n = 54 and 114 from seven embryos for 2-APB treatment. (D, E) Formation of c-lamellipodia by surrounding cells inside and outside the Ca^2+^ wave. The frequency (D) and size (E) of c-lamellipodia were determined. n = 115 and 33 from five embryos for control, n = 21 and 38 from seven embryos for Gd^3+^ treatment, n = 68 and 68 from ten embryos for GsMTx4 treatment, n = 31 and 33 from six embryos for XestoC treatment, and n = 52 and 67 from six embryos for 2-APB treatment. Error bars show mean± SD. n.s., not significant; \**P* < 0.05, and \*\**P* < 0.01 (paired t-test).

**Figure 4.**
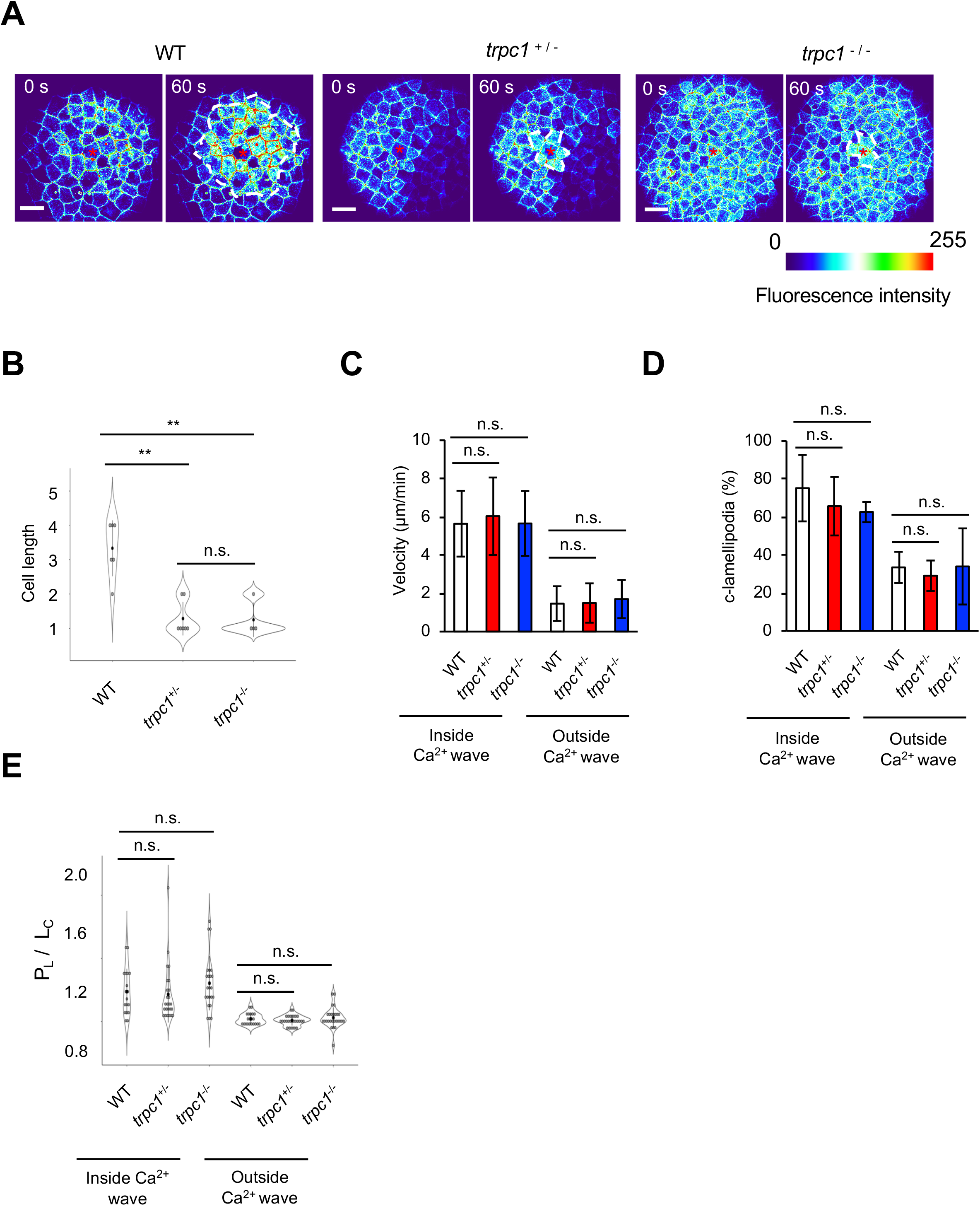
*trpc1* is involved in Ca^2+^ wave propagation. (A) Color-coded images of GCaMP7 and Lifeact-GFP in WT, heterozygous (*trpc1*^+/-^), and homozygous (*trpc1*^-/-^) embryos at 0 and 60 s after laser irradiation. White dotted lines indicate the area of Ca^2+^ wave propagation. Red asterisks indicate the laser focal point. Scale bar: 50 μm. (B–E) Effects of *trpc1* knockout on Ca^2+^ wave-mediated polarized movements. (B) The expansion area of the Ca^2+^ wave. n = 6, 7, and 4 embryos. (C) Polarized movements of surrounding cells inside and outside the Ca^2+^ wave. (D, E) Formation of c-lamellipodia by surrounding cells inside and outside the Ca^2+^ wave. The frequency (D) and size (E) of c-lamellipodia were determined. n = 27 and 36 from six WT embryos, 45 and 21 from nine *trpc1^+/-^* embryos, and n = 45 and 21 from three *trpc1*^-/-^ embryos. (E) n = 18 and 21 from six WT embryos, 25 and 28 from nine *trpc1^+/-^* embryos, and n = 23 and 27 from three *trpc1^-/-^* embryos. Error bars show mean±SD. n.s., not significant and \*\**P* < 0.01 (paired t-test).

### The Ca^2+^ wave promotes the polarized movements of surrounding cells through the formation of c-lamellipodia in zebrafish

The Ca^2+^ wave regulates the collective movements of cells (*15–17*). Mechanosensitive Ca^2+^ channels promote Ca^2+^ entry and induce release of Ca^2+^ from the ER through the IP_3_ receptor via a process called store-operated calcium entry (SOCE) (*18–21*). Thus, we investigated the roles of Ca^2+^ influx, mediated by mechanosensitive Ca^2+^ channels, and Ca^2+^ release from the ER through IP_3_ receptors in Ca^2+^ wave-mediated polarized movements during cell extrusion in zebrafish. In embryos treated with Gd^3+^ or GsMTx4 (mechanosensitive Ca^2+^ channel inhibitors) and *trpc1* mutants, the Ca^2+^ wave was not propagated by the second/third or more outer rows of surrounding cells (distal cells), and polarized movements and c-lamellipodia formation in distal cells were diminished (Fig. 3A, C–E, Fig. 4), suggesting that Ca^2+^ influx mediated by mechanosensitive Ca^2+^ channels regulates Ca^2+^ wave propagation toward distal cells. However, in proximal cells with elevated Ca^2+^, which were located in the first and second rows of surrounding cells, polarized movements and c-lamellipodia formation occurred normally (Fig. 3C–E and Fig. 4C–E), suggesting that Ca^2+^ elevation in the cytoplasm mediated by SOCE is required for polarized movements and c-lamellipodia formation. Similar to the effects of mechanosensitive Ca^2+^ channel inhibition, treatment with 2-APB or XestoC (IP_3_ receptor inhibitors) inhibited Ca^2+^ wave propagation toward distal cells (Fig. 3A, B), suggesting that IP_3_ receptor-mediated Ca^2+^ release from the ER is involved in Ca^2+^ wave propagation toward distal cells. However, inconsistent with the effects of mechanosensitive Ca^2+^ channel inhibition, inhibition of the IP_3_ receptor significantly suppressed polarized movements and c-lamellipodia formation in proximal cells, in which the Ca^2+^ level was elevated (Fig. 3C–E). These results suggest that the IP_3_ receptor is involved in Ca^2+^ wave propagation, polarized movements, and formation of c-lamellipodia and that SOCE mediated by the IP_3_ receptor is required for polarized movements through the formation of c-lamellipodia during cell extrusion.

### Difference between Ca^2+^ wave-mediated polarized movements and actomyosin ring contraction

Ca^2+^ wave-mediated polarized movements occurred before contraction of the actomyosin ring (Fig. 1A and Video S1). To investigate whether Ca^2+^ wave-mediated polarized movements are an initial or independent process of actomyosin ring contraction, we ablated a single contact edge between the extruding and surrounding cells using a laser. During actomyosin ring contraction, the contact edge becomes short (*22*). Therefore, when the contact edge was cut by the laser during actomyosin ring contraction (approximately 120 s after laser irradiation), the distance between the vertexes of the contact edge greatly increased (Fig. 5A, C, D), indicating that actomyosin ring contraction generates contractile tension during cell extrusion, as shown previously (*22*). By contrast, when the contact edge was cut during Ca^2+^ wave-mediated polarized movements (approximately 30 s after laser irradiation), the distance between the vertexes of the contact edge did not change at all (Fig. 5B, D), indicating that contractile tension is not loaded on the edge during Ca^2+^ wave-mediated polarized movements. These data suggest that Ca^2+^ wave-mediated polarized movements are independent of actomyosin ring contraction.

**Figure 5.**
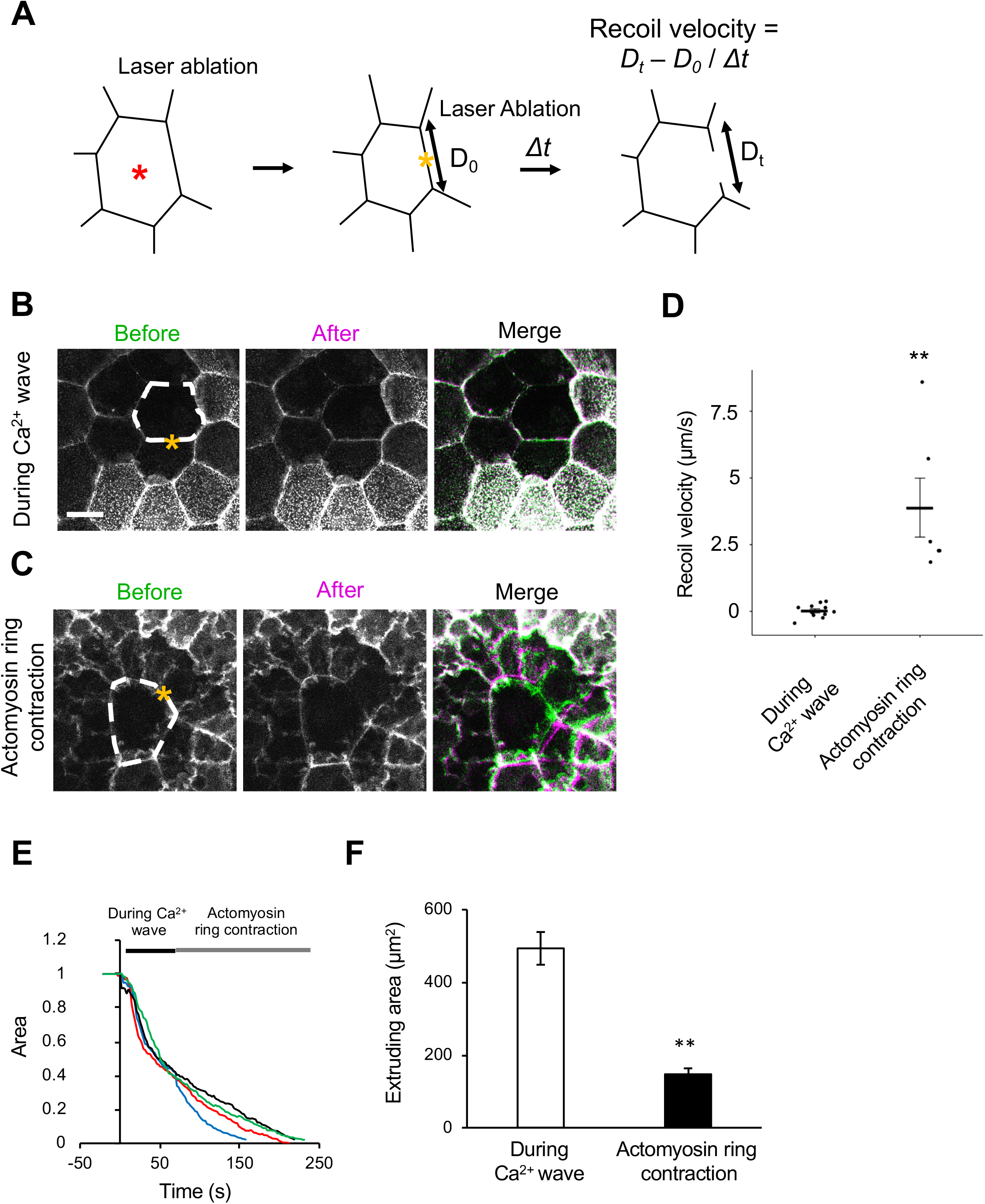
Mechanical properties of surrounding cells during cell extrusion. (A) Schematic diagram of measurement of the contact edge recoil velocity following laser ablation of a single contact edge between extruding and surrounding cells. An extruding cell is induced by laser irradiation of the cell center (red asterisk). A single contact edge between the extruding cell and a surrounding cell is cut by laser ablation (yellow asterisk). The recoil velocities (*D_t_* – *D*_0_ / *Δt*) were evaluated from the initial edge distance (*D*_0_) and recoiled distance (*D_t_*) at a time after laser cutting (*Δt*). (B–D) Fluorescence images of recoil behaviors during the Ca^2+^ wave and actomyosin ring contraction. Representative images of recoil behaviors during the Ca^2+^ wave (B) and actomyosin ring contraction (C). Fluorescence images of Lifeact-GFP before (green) and after (purple) laser cutting were merged. Scale bar: 10 μm. (D) Recoil velocities during the Ca^2+^ wave and actomyosin ring contraction. n = 11 and 6 embryos. (E) Reduction of the area of extruding cells through cell extrusion. Data are from four embryos. The black and gray bars at the top indicate the period of Ca^2+^ propagation and actomyosin ring contraction, respectively. (F) The reduction rate of the area of extruding cells during the Ca^2+^ wave (white) and actomyosin ring contraction (black) were measured from (E). Data are mean ± SD. \*\**P* < 0.01 (paired t-test).

### Force generation during Ca^2+^ wave-mediated polarized movements

The area of extruding cells decreased by 61.3 ± 7.4% during the Ca^2+^ wave (by approximately 60 s after laser irradiation) (Fig. 5E, F and Table 1). Thus, we assumed that this process generates a force to fill the space immediately. To test this possibility, we used the *in vivo* force measurement method that we previously established and measure the force generated in this process. First, the center of a cell was irradiated with a single pulse of a femtosecond laser to initiate cell extrusion (Fig. 6A, left panel). Next, when surrounding cells had collectively moved toward the extruding cell (approximately 30 s after laser irradiation), a series of impulsive forces was loaded (44 times at 1 s intervals) on the center of the extruding cell (Fig. 6A, right panel). Consequently, the polarized movements were counter-balanced by the force loading (Fig. 6B, C). Thereafter, when force loading was stopped, the polarized movements restarted (Fig. 6B, C, and Video S2).

**Table 1.** Force and reduction rate of the extruding cell area during Ca^2+^-mediated polarized movements.

|  | Pressure (kPa) | Reduction rate (%) |
| --- | --- | --- |
| Control | 1.08 ± 0.17 (n = 33) | 61.3 ± 7.4 (n = 17) |
| Gd<sup>3+</sup> | 0.97 ± 0.08 (n = 19)** | 49.8 ± 4.0 (n = 5)** |
| GsMTx4 | 0.86 ± 0.13 (n = 16)** | 47.5 ± 5.7 (n = 9)** |
| XestoC | 0.99 ± 0.13 (n = 22)* | 31.8 ± 11.7 (n = 7)** |
| 2-APB | 0.94 ± 0.17 (n = 33)** | 23.1 ± 11.7 (n = 8)** |
| *trpc1*<sup>-/-</sup> | 0.82 ± 0.14 (n = 21)* | 44.0 ± 7.1 (n = 10)** |

n indicates the number of embryos.
\**P* < 0.05 and \*\**P* < 0.01 (paired t-test).

**Figure 6.**
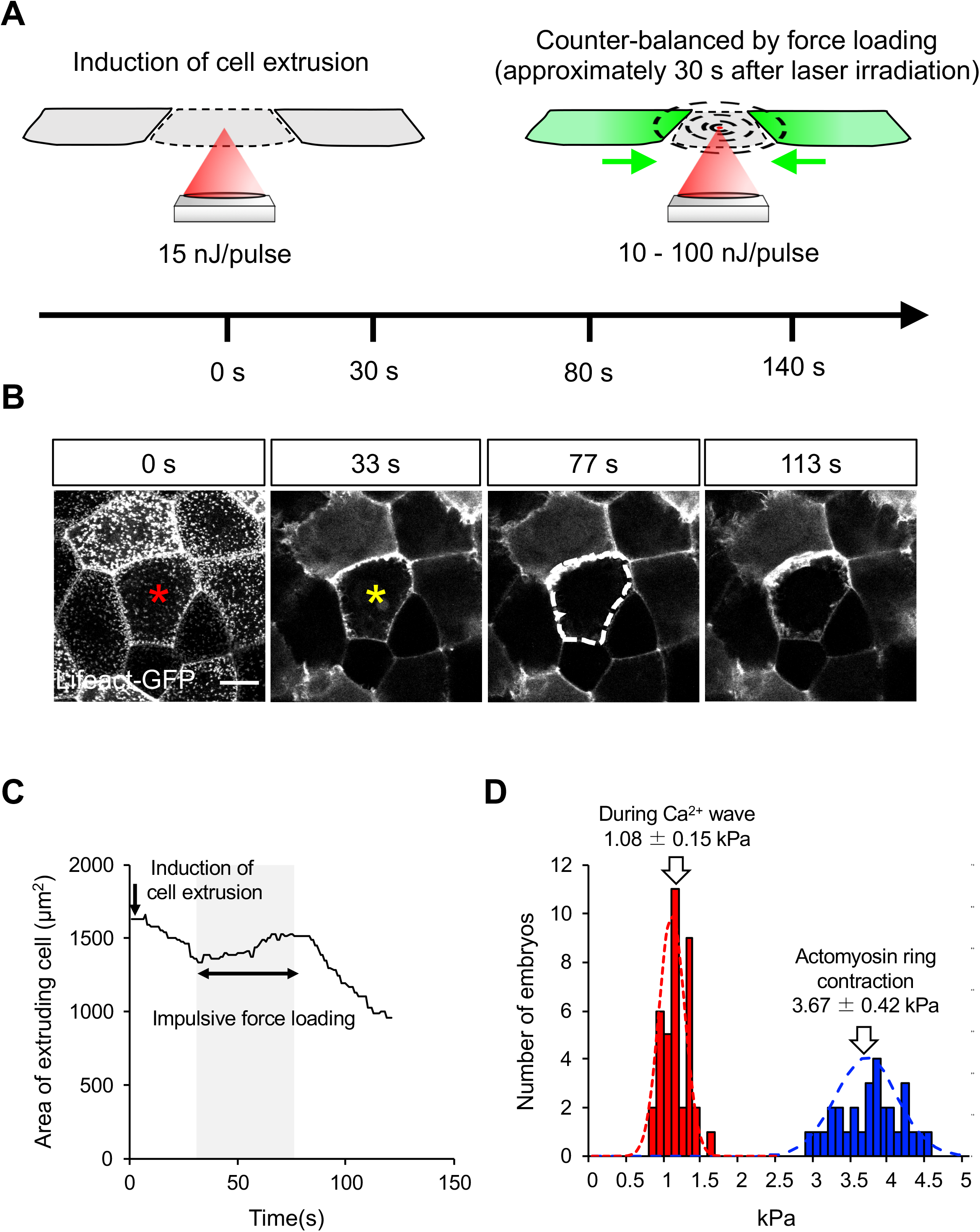
Ca^2+^ wave-mediated polarized movements generate approximately 1 kPa of force. (A) Schematic illustrations of force measurement using a femtosecond laser. Left panel, cell extrusion is induced by irradiation of the cell center with a femtosecond laser (15 nJ/pulse). Right panel, after Ca^2+^ wave-mediated polarized movements are initiated (approximately 30 s after laser irradiation), a series of laser pulses (10–100 nJ/pulse) is loaded every 1 s at the center of the extruding cell. (B, C) Force measurement during Ca^2+^ wave-mediated polarized movements. (B) Images were extracted from Video S2. Red and yellow asterisks indicate the focal point of the laser to induce the extruding cell and loading of impulsive forces, respectively. The white dotted line at 77 s shows the size of the extruding cell at 39 s. (C) Plot of the area of the extruding cell over time. An extruding cell was induced by laser irradiation at 0 s and gradually became smaller. By loading impulsive force, the size of the extruding cell slightly increased. After stopping the loading force, the extruding cell became smaller over time. (D) Estimated forces generated by Ca^2+^-mediated polarized movements (red, n = 34) and actomyosin ring contraction (blue, n = 29). The mean value and SD of the pressure are also indicated.

The impulsive force propagated spherically as a volume wave from the vicinity of the laser focal point; therefore, the force for counter-balance was estimated from the pressure (*P*) acting at distance (*R*) (Fig. S1A, see also Materials and Methods) (*9*). To investigate the statistical accuracy of the measurement, we evaluated the *P* calculated by Eq. [2] and summarized each data point in Figure S1B as a histogram (Fig. 6D). The data had a Gaussian distribution, and the mean value and SD were reliable compared with the *P* estimated in Figure S1B. Based on these correlations, the *P* of the Ca^2+^-mediated polarized movements was estimated to be 1.08 ± 0.17 kPa (Table 1). This value is almost 4 times smaller than the force of actomyosin ring contraction (Fig. 6D) (*9*). Taken together, our results demonstrate that relatively small forces generated by two distinct processes (i.e., Ca^2+^-mediated polarized movements and actomyosin ring contraction) drive cell extrusion in zebrafish.

Finally, we calculated the *P* [Ca^2+^-mediated polarized movements] in embryos treated with several inhibitors and in *trpc1* homozygous mutants to determine which molecules contribute to force generation during Ca^2+^ wave-mediated polarized movements. To investigate the relationship between the *P* and mechanosensitive Ca^2+^ channels, we measured the force in embryos treated with Gd^3+^ or GsMTx4 and in *trpc1* homozygous mutants (Fig.7, S2). Compared with controls, both the *P* and reduction rate of the extruding cell area during the Ca^2+^ wave were significantly reduced in these experimental settings, suggesting that mechanosensitive Ca^2+^ channels including trpc1 contribute to force generation during cell extrusion. Similar results were obtained when we inhibited the IP_3_ receptor using XestoC or 2-APB. Although the *P* did not significantly differ between samples in which mechanosensitive Ca^2+^ channels and the IP_3_ receptor were inhibited, the reduction rate of the extruding cell area was smaller upon inhibition of the IP_3_ receptor than upon inhibition of mechanosensitive Ca^2+^ channels (Table 2). Since inhibition of the IP_3_ receptor, but not mechanosensitive Ca^2+^ channels, blocked polarized movements through c-lamellipodia formation (Fig.3, 4), we suggest that the c-lamellipodia formation is involved in the reduction rate of the extruding cell area during the Ca^2+^ wave.

**Figure 7.**
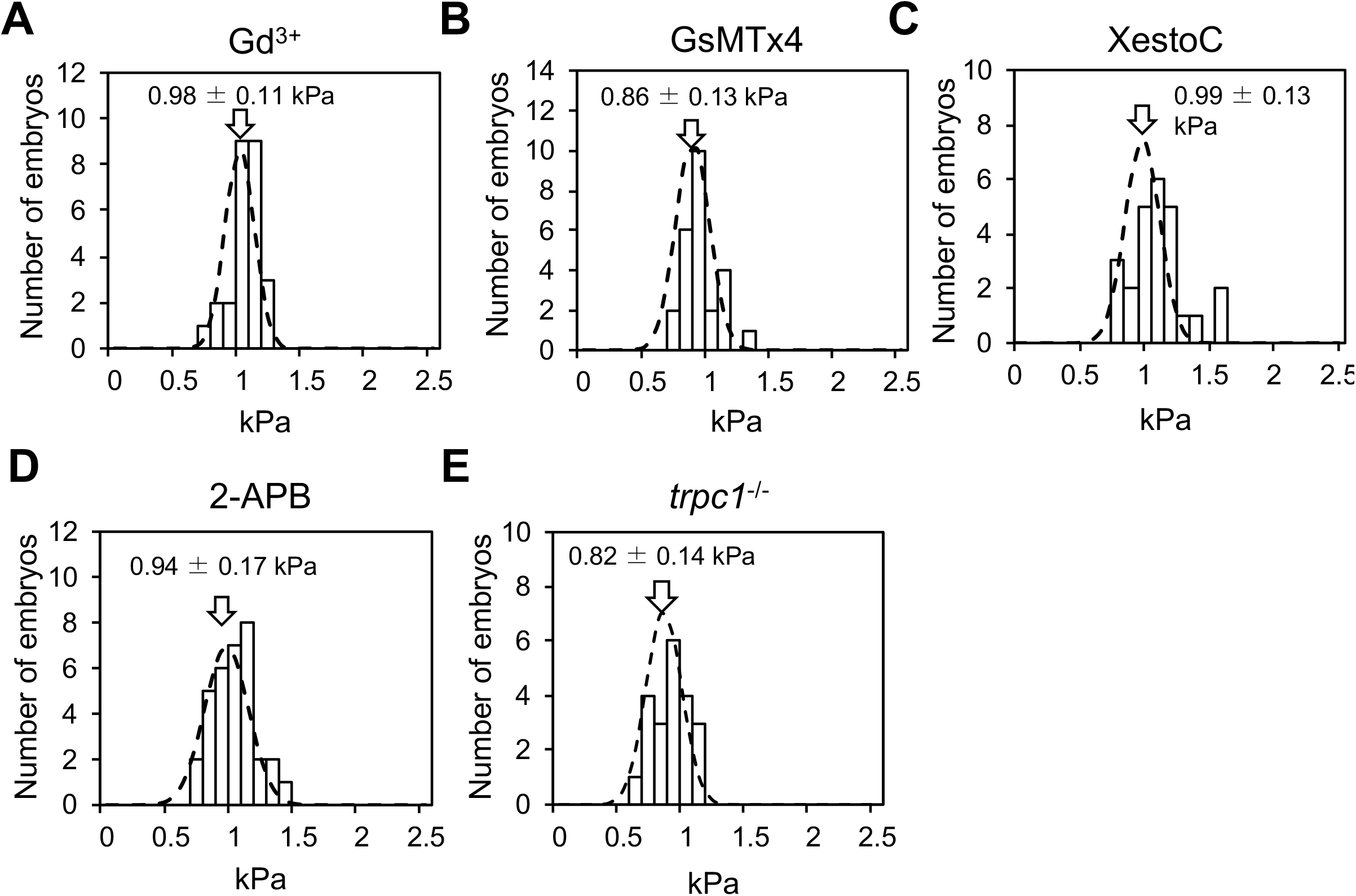
Effects of mechanosensitive Ca^2+^ channels and the IP_3_ receptor on forces generated by Ca^2+^ wave-mediated polarized movements. (A-E) Forces generated by Ca^2+^ mediated-polarized movements were estimated in embryos treated with Gd^3+^ (n = 19, A), GsMTx4 (n = 16, B), XestoC (n = 22, C), and 2-APB (n = 33, D), and in *trpc1^-/-^* embryos (n = 21, E). See also Table 1. The mean value and SD of the pressure are also indicated.

**Table 2.**
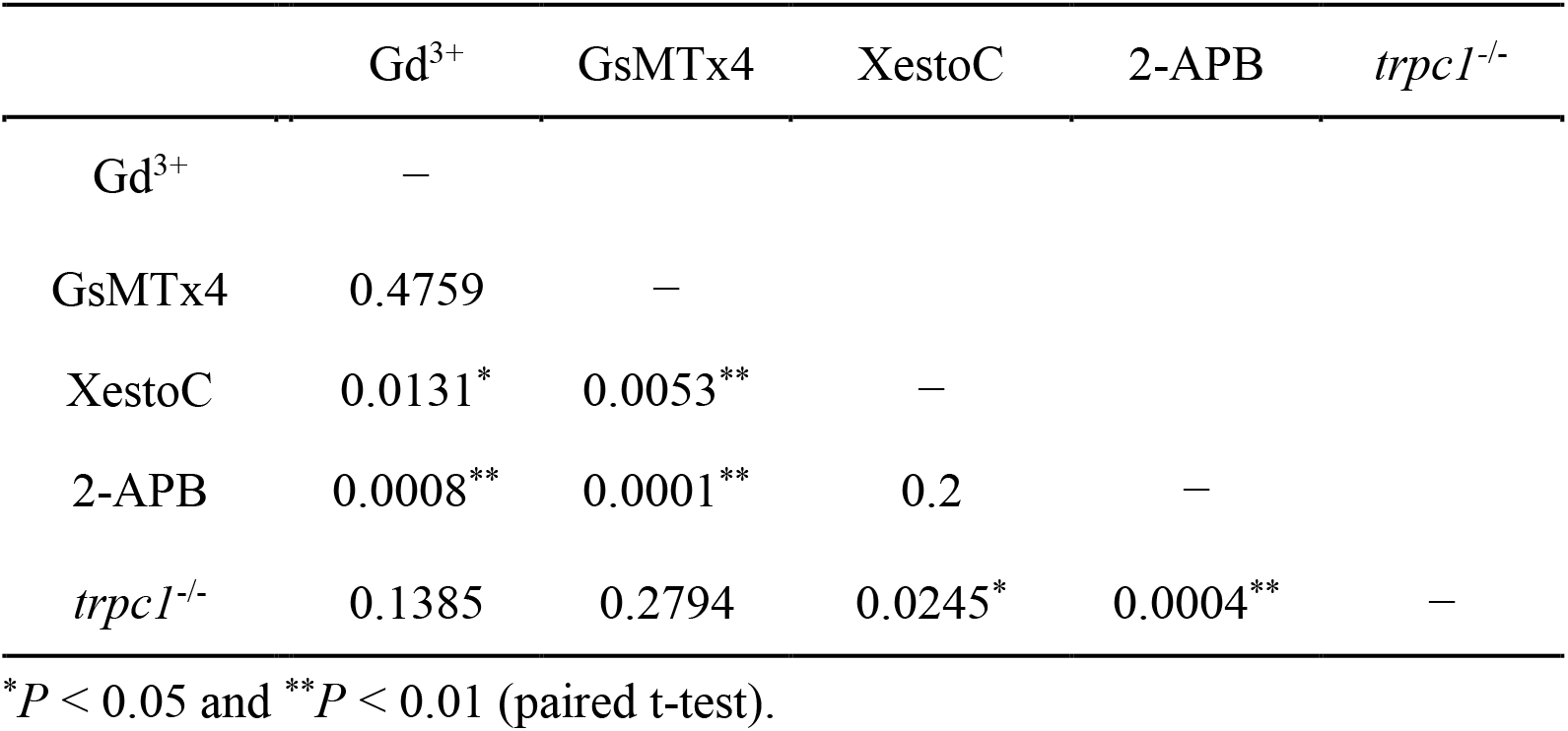
Statistical comparison of the reduction rate of the extruding cell area upon inhibition of mechanosensitive Ca^2+^ channels and the IP_3_ receptor.

## Discussion

A Ca^2+^ wave regulates cell extrusion in mammalian cell cultures and zebrafish (*10*). In this study, we characterized its molecular mechanism in zebrafish and measured the force it generates. We demonstrated that mechanosensitive Ca^2+^ channel-dependent Ca^2+^ entry and IP_3_ receptor-dependent Ca^2+^ release initiate the Ca^2+^ wave and propagate it toward multiple rows of surrounding cells and that the IP_3_ receptor is required for actin rearrangements to form c-lamellipodia, which facilitate the collective movements of surrounding cells toward extruding cells (Fig. 8). Based on force measurement, we propose that polarized movements of surrounding cells through c-lamellipodia formation help to generate 1 kPa of force during Ca^2+^ wave-mediated cell extrusion (Fig. 6). We previously demonstrated that actomyosin ring contraction generates force during cell extrusion in zebrafish and estimated that the *P* is approximately 4 kPa (*9*). Here, we revealed that Ca^2+^ wave-mediated polarized movements are one of the force-generating processes during cell extrusion and occur before actomyosin ring contraction, and that this force is smaller than the contractile tension generated by actomyosin ring contraction.

**Figure 8.**
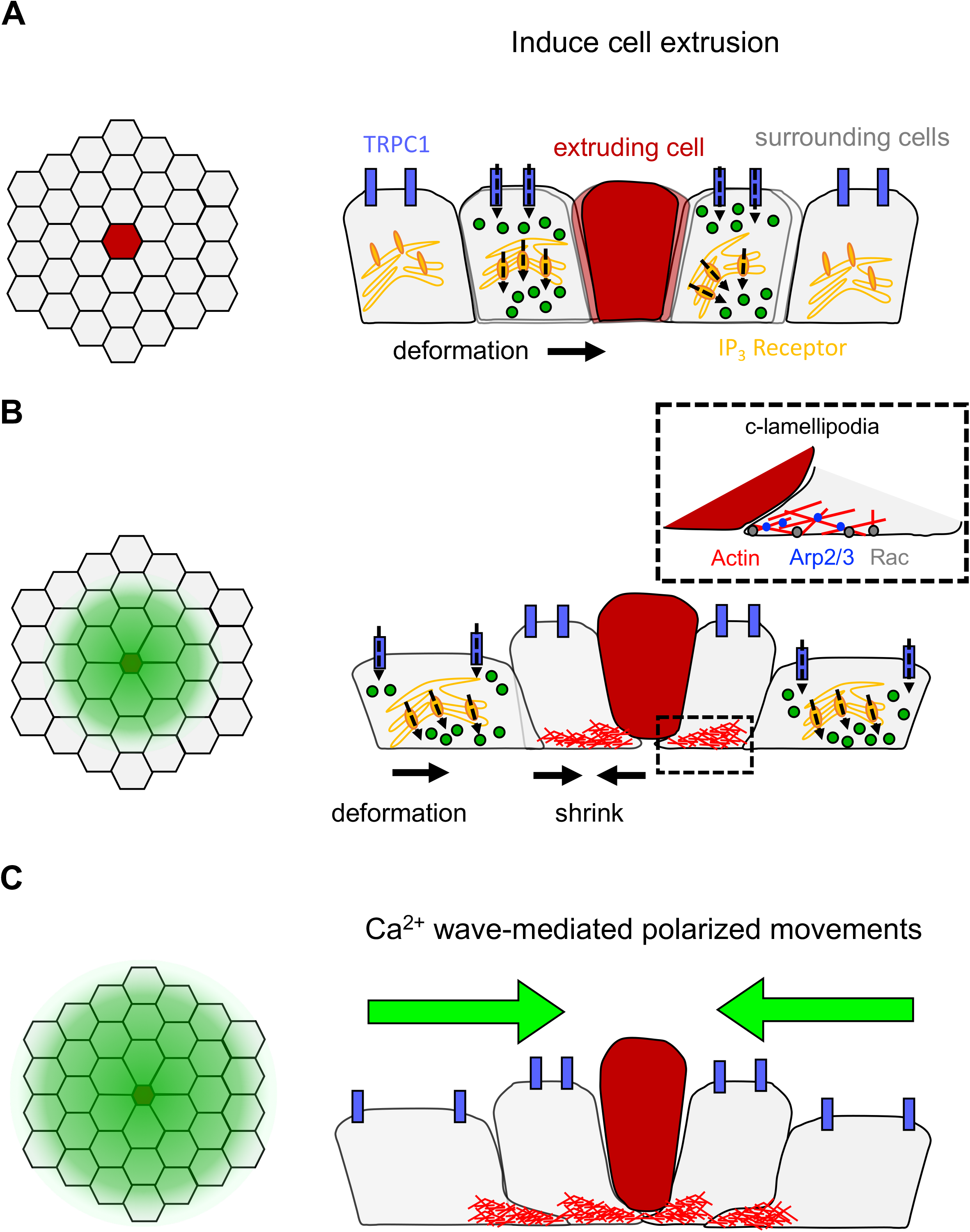
Cell extrusion in zebrafish embryos. Schematic representation of cell extrusion in zebrafish proposed in this study. (A) Cell extrusion leads to deformation of cells surrounding the extruding cell. Cell deformation induces an increase in intracellular Ca^2+^ through TRPC1 and SOCE, resulting in cell shrinkage. (B) This shrinkage induces deformation of cells located in the second row of surrounding cells. The increase of intracellular Ca^2+^ leads to actin rearrangement in the cells, resulting in c-lamellipodia formation. (C) This chain of reactions occurs in several rows of surrounding cells, leading to Ca^2+^-mediated polarized movements.

Mechanosensitive Ca^2+^ channels including TRPC1 are activated by membrane stretching and SOCE (*19–21*). IP_3_, which is propagated through gap junctions, activates the IP_3_ receptor, which contributes to SOCE (release of Ca^2+^ from the ER) (*10*). In mammalian cell cultures, inhibition of mechanosensitive Ca^2+^ channels, the IP_3_ receptor, and gap junctions blocks propagation of the Ca^2+^ wave toward distal surrounding cells (*10*). Thus, both SOCE via TRPC1 and the IP_3_ receptor and IP_3_ propagation through gap junctions are involved in Ca^2+^ wave propagation. However, pharmacological experiments showed that mechanosensitive Ca^2+^ channels including trpc1 and IP_3_ receptor, but not gap junctions, are involved in Ca^2+^ wave propagation in zebrafish embryos, suggesting that that the Ca^2+^ wave propagates without gap junctions in zebrafish. We assume that a chain reaction of mechanotransduction from proximal to distal cells through mechanosensitive Ca^2+^ channels regulates this process. When cell extrusion is initiated, the first row of surrounding cells is deformed, inducing activation of mechanosensitive Ca^2+^ channels including trpc1. Dependent on entry of a small amount of Ca^2+^, SOCE is activated in a cell, resulting in its shrinkage. This induces stretching of a cell located in the second row of surrounding cells. These chain reactions occur in several rows of surrounding cells, but mechanotransduction gradually decreases and stops the propagation. This idea is supported by the findings that when the first row of surrounding cells translocated toward the extruding cell during the Ca^2+^ wave, they slightly deformed to fill 60% of the area occupied by the extruding cell (Fig. 5F and Table 1) and that Ca^2+^ elevation induced shrinkage of a cell within an epithelial sheet, which provoked stretch-dependent Ca^2+^ entry in adjacent cells. Additional experiments are required to elucidate how the Ca^2+^ wave propagates from a cell to other cells and how propagation stops.

Zebrafish *trpc1* encodes a protein containing 796 aa and six transmembrane domains. A tetramer of trpc1 forms the mechanosensitive Ca^2+^ channel. In the *trpc1* knockout mutants generated in this study, 13 bp are deleted in exon 4 of the *trpc1* gene and the mutated *trpc1* encodes a truncated protein containing only 168 aa, which lacks almost all functional domains of trpc1 (Fig. S3). Consequently, we think this mutant is a knockout. Although the Ca^2+^ wave failed to propagate in *trpc1* heterozygous and homozygous zebrafish mutants, we do not observe any phenotypical differences between these two mutants. Based on the failure of the Ca^2+^ wave to propagate in heterozygous mutants, the mutation may have a dominant-negative effect on Ca^2+^ wave propagation. However, two lines of evidence suggest this is unlikely. First, Ca^2+^ wave propagation was significantly inhibited in mammalian cultured cells following shRNA-based knockdown of TRPC1, although the knockdown efficiency was approximately 50% (*10*). Second, we observed cardiac failure in *trpc1* homozygous embryos, but not in heterozygous *trpc1* embryos (manuscript in preparation). These findings suggest that reduction of the amount of trpc1 protein in the mutants leads to a functional defect of mechanosensitive Ca^2+^ channels and that the amount of trpc1 protein is critical for Ca^2+^ wave propagation during cell extrusion.

By using femtosecond laser impulsive forces, we estimated that approximately 1 kPa of force is generated by Ca^2+^ wave-mediated polarized movements; this is lower than contractile tension (4 kPa) generated by actomyosin ring contraction during cell extrusion. However, we think approximately 1 kPa is a lower limit of this quantification method because we observed a significant difference in the reduction rate of the extruding cell area, but not in force, between samples in which mechanosensitive Ca^2+^ channels or the IP_3_ receptor were inhibited (Tables 1 and 2). Our force quantification method *in vivo* is useful for force estimation of physiological processes in living organisms, but we must establish a new method to improve its quantitativeness in the future.

The Ca^2+^ wave is the evolutionarily conserved and general regulatory mechanism of cell extrusion. In this study, we characterized the molecular mechanisms of the Ca^2+^ wave in zebrafish and demonstrated that the detailed mechanisms differ between zebrafish embryos and mammalian cultured cells, whereas cellular behaviors are conserved. Furthermore, we revealed that Ca^2+^ wave-mediated polarized movements are one of the force-generating processes during cell extrusion.

## Materials and Methods

### Zebrafish experiments

WT zebrafish and *Tg[krt4:Lifeact-GFP]* lines were used in this study. All zebrafish experiments were performed with the approval of the Animal Studies Committee of the Nara Institute of Science and Technology.

### mRNA synthesis and injection

pCS2 carrying *Lifeact-GFP* (a gift from Dr. Noriyuki Kinoshita), *GCaMP7* (a gift from Dr. Junichi Nakai), *RFP*, or *RacN17* was used as a template for mRNA synthesis. mRNAs were synthesized using the SP6 mMessage mMachine System (Thermo Fisher Scientific). The injection was performed as described previously (*29*). To observe both F-actin and Ca^2+^ dynamics, *Lifeact-GFP* mRNA (100 pg) and *GCaMP7* mRNA (100 pg) were co-injected into WT embryos at the one-cell stage or *GCaMP7* mRNA (100 pg) was injected into embryos at the one-cell stage obtained from the *Tg[krt4:Lifeact-GFP]* line.

### Treatment with inhibitors

Embryos were developed until 5 hours post-fertilization (hpf), treated with 100 μM Gd^3+^ (439770; Sigma-Aldrich), 4 μM GsMTx4 (095-05831; FUJIFILM WAKO Pure Chemical Corporation), 4 μM XestoC (244-00721; FUJIFILM WAKO Pure Chemical Corporation), 50 μM 2-APB (D9754; Sigma-Aldrich), 100 μM CK666 (SML0006; Sigma-Aldrich), 10 μM Y27632 (08945-84; Nacalai Tesque), or 50 μM Blebbistatin (856925-71-8; Sigma-Aldrich) for 60 min and then used in the following experiments. Non-treated or DMSO (vehicle)-treated embryos were used as negative controls.

### Observation of F-actin and Ca^2+^ dynamics during cell extrusion in zebrafish

Embryos expressing Lifeact-GFP and GCaMP7 were developed until around 6 hpf, dechorionated, and mounted in the holes of a gel made with 1% low-melting point agarose (Nacalai Tesque) on 35 mm glass-bottom dishes (Matsunami). A single pulse of a femtosecond laser (15 nJ/pulse; Solstice Ace, Spectra-Physics) was focused through a 100×/1.25 objective lens (Olympus) onto the centers of epithelial cells located near the animal poles of embryos at 6 hpf, as described previously (*9*). F-actin and Ca^2+^ dynamics were observed for 1–7 min at 1–15 s intervals. At each time point, Z-stack images of the embryos (8–17 planes at 1 or 2 μm intervals) were also obtained.

### CRISPR-based knockout of *trpc1* in zebrafish

Guide sequences of *trpc1* single-guide RNA targeting zebrafish *trpc1* were designed based on exon 4 using the CRISPRdirect web server. The synthesized cgRNA for *trpc1* (5’-UCCAUGGUGGUGGAAUACUCguuuuagagcuaugcuguuuug-3’) and tracrRNA (5’ - AAACAGCAUAGCAAGUUAAAAUAAGGCUAGUCCGUUAUCAACUUGAAAAAGUGGCACCGAGUCGGUGCU-3’), both of which act as the single-guide RNA, were obtained from Fasmac. The cgRNA (100 pg), tracrRNA (100 pg), and Cas9 protein (500 pg) (GE-005; Fasmac) were co-injected into zebrafish embryos at the one-cell stage. By screening germline-transmitted lines, a line carrying the 13 bp deletion in exon 4 (Fig. S3) was obtained and the F2 generations were used in this study. Genotyping was performed by PCR using sets of site-specific primers for the WT (*trpc1-WT*-L, 5’-TACAGAAGATTCAGAACCCAGAGTATT-3’; and *trpc1-WT*-R, 5’-CAGATCCATTTGTTCTCCTTCTATGA-3’) and mutant (*trpc1-13del*-L, 5’-TACAGAAGATTCAGAACCCACCA-3’; and *trpc1-WT*-R) alleles.

### Quantification of mechanical force in zebrafish

A single pulse of a femtosecond laser (15 nJ/pulse) was focused onto the center of an epithelial cell. Next, when polarized movements started toward the extruding cell (approximately 30 s after laser irradiation), a series of impulsive forces (10–100 nJ/pulse) was loaded (40-50 times at 1 s intervals) at the center of the extruding cell. Dynamic changes of F-actin were observed for 5–7 min at 1 s intervals. The area of the extruding cell (μm^2^) at each time point was measured by ImageJ software (NIH). In addition, the counter-balanced radius *R* (μm) of the extruding cell was measured. Using the measurement results and Eq. [1]–[2], the force generated in this process was estimated as described in the Results.

### Quantification of impulsive force by atomic force microscopy

When the femtosecond laser is focused in the vicinity of an atomic force microscope cantilever, the total force *F*_0_ generated at the laser focal point is estimated from the bending movement of the cantilever. The details have been described previously (*9*). Based on this estimation, the impulsive force *F*_0_ generated at the laser focal point is related to the incident laser pulse energy *L* by the following equation:

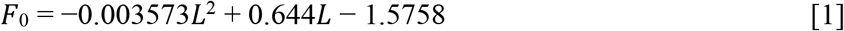

Assuming that *F*_0_ propagates spherically as a volume wave in the vicinity of the laser focal point, the impulsive force *F*_0_ generated at the laser focal point as a unit of pressure *P* is expressed by the following equation:

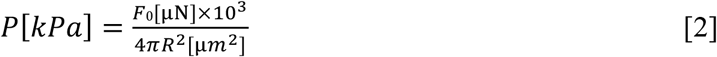

In Figure S1, the vertical and horizontal axes of Figure 1B have been respectively converted from *R* to *R*^2^ and from *L* to *F*_0_. The relationship is almost linear (*R*^2^ = 0.8); therefore, the pressure is estimated from the slope of the least-squares fitting line by Eq. [2].

### Statistical analysis

Differences in means were analyzed by the paired t-test. The results of the t-test were considered significant when *P* < 0.05.

## Acknowledgments

We thank Ryohei Yasukuni and Kazunori Okano for fruitful discussions. We are also grateful to Noriyuki Kinoshita and Junichi Nakai for sharing *Lifeact-GFP* and *GCaMP7*, respectively, and to Maiko Yokouchi and Ayasa Kuroda for technical assistance with zebrafish experiments.

## Funding

This work was supported by ACT-X Grant Number JPMJAX191K (S.Y.); JSPS KAKENHI Grant Numbers JP19H04782, JP17H05768, JP18H02451 (Y.B.), and JP17H05621 (T.M.); the Takeda Science Foundation and Suntory Foundation for Life Sciences (T.M.); the SUMBOR grant (T.M.); the Foundation of Nara Institute of Science and Technology (S.Y. and T.M.); and AMED under Grant Number 21gm0810011h0005 (Y.H.).

## Author contributions

S.Y. performed almost all experiments and data analysis. Y.H. calibrated the femtosecond laser-induced impulsive force and supervised the femtosecond laser analysis. Y.B., Y.F., Y.H., and T.M. conceived the project. S.Y., Y.H., and T.M. designed the experiments and wrote the manuscript. All authors reviewed the manuscript.

## Notes

### Competing Interest Statement

The authors have declared no competing interest.

