## Supplementary Information for "A Ca^2+^ wave generates a force during cell extrusion in zebrafish"

### Supplementary Figures

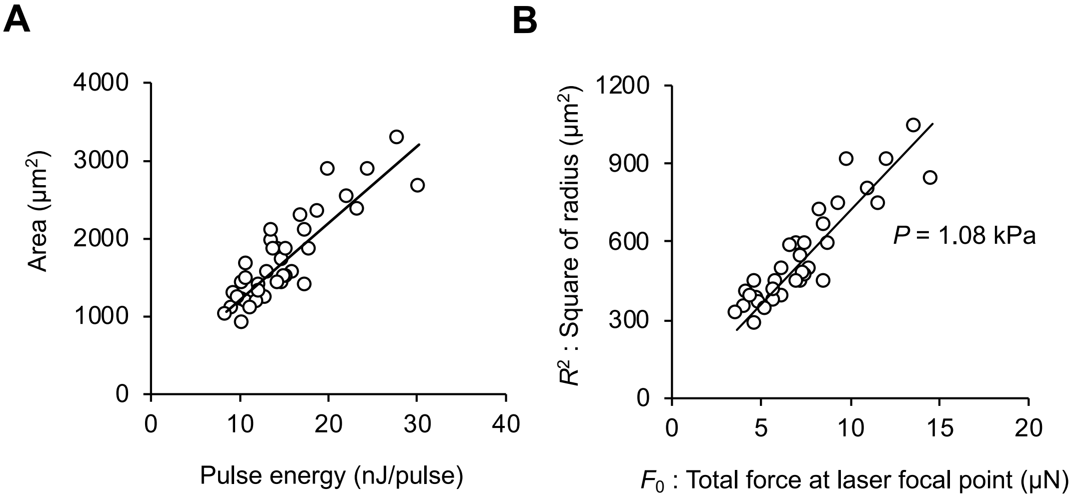


**Supplementary Figure 1. Measurement of mechanical force required for cell extrusion in controls.** (A) The area of the extruding cell in controls was plotted against laser pulse energy (n = 34), revealing a linear positive correlation (correlation coefficient, $R^2 = 0.77$). (B) The square of the radius of the extruding cell was plotted against total force at the laser focal point $F_0$ (n = 34), revealing a linear positive correlation (correlation coefficient, $R^2 = 0.8$).

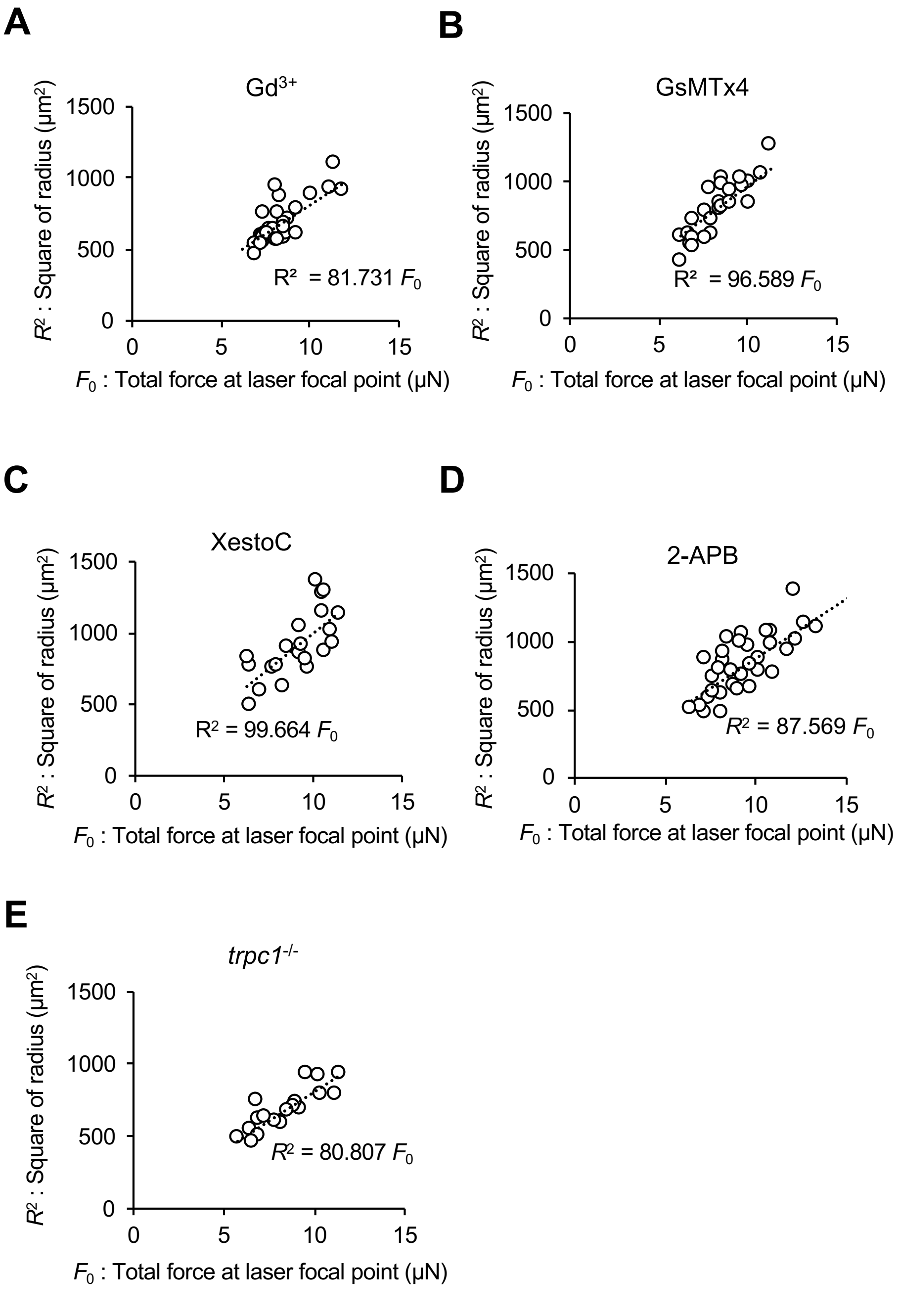


**Supplementary Figure 2. Measurement of force in manipulated embryos.** The square of the radius $R^2$ is proportional to total force $F_0$ in embryos treated with $Gd^{3+}$ (A), GsMTx4 (B), XestoC (C), and 2-APB (D) and in *trpc1*$^{-/-}$ embryos (E) (correlation coefficient, $R^2 = 0.61$, 0.71, 0.51, 0.54, and 0.53). The magnitude of the force in these experimental settings was estimated and is listed in Table 1.

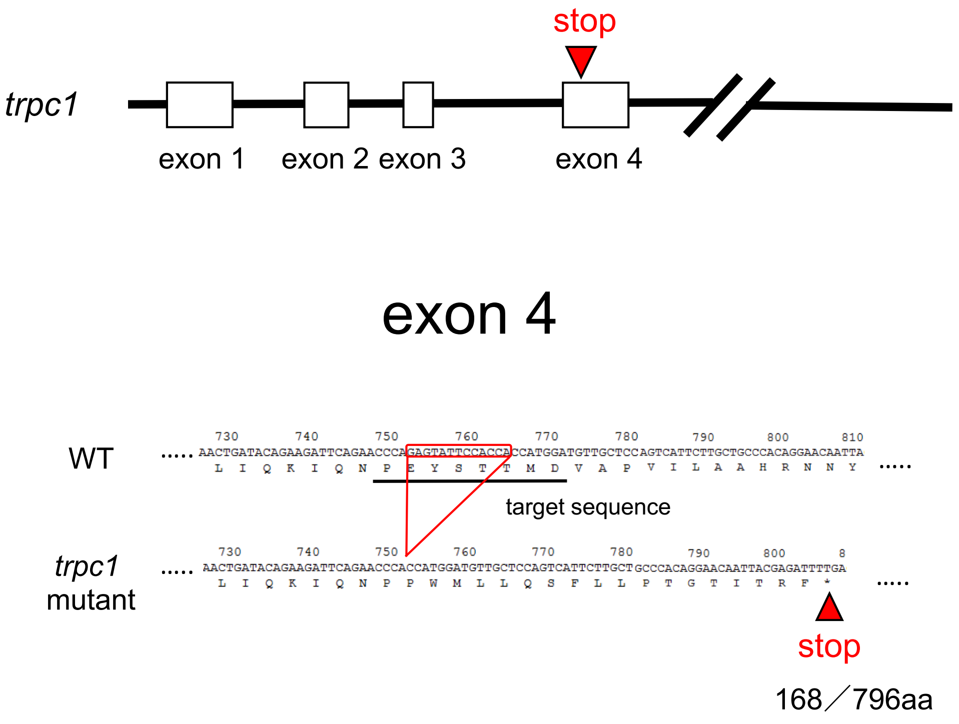


**Supplementary Figure 3. Schematic of the *trpc1* mutant generated using CRISPR.** Mutation sites in the *trpc1* mutant generated using CRISPR. Red triangles indicate the insertion sites of the STOP codon.

### Supplementary Video Legends

**Supplementary Video 1. $Ca^{2+}$ wave propagation during cell extrusion in zebrafish**

The center of an epithelial cell in the GCaMP7-expressing *Tg[krt4:Lifeact-GFP]* line was directly irradiated with a single pulse of a femtosecond laser (15 nJ/pulse) at time = 0 s. Images were taken every 2.7 s. Scale bar: 50 μm.

**Supplementary Video 2. Force measurement experiment**

The center of an epithelial cell in a Lifeact-GFP-expressing embryo was directly irradiated with a single pulse of a femtosecond laser (15 nJ/pulse) at time = 0 s. After polarized movements started at around 39 s, impulsive forces generated by the femtosecond laser (25 nJ/pulse) were loaded onto the center of the actomyosin ring 40 times at 1 s intervals (red dots at 39–77 s). Images were taken every 1 s. Scale bar: 10 μm.
